# Overlapping connectivity gradients in the anterior temporal lobe underlie semantic cognition

**DOI:** 10.1101/2020.05.28.121137

**Authors:** Myrthe Faber, Izabela Przeździk, Guillén Fernández, Koen V. Haak, Christian F. Beckmann

**Author notes:** These authors contributed equally as co-first authors (Myrthe Faber, Izabela Przeździk). These authors contributed equally as co-last authors (Koen V. Haak, Christian F. Beckmann). **Corresponding author:** Myrthe Faber, Donders Institute for Brain, Cognition and Behaviour, Donders Centre for Cognitive Neuroimaging, Kapittelweg 29, 6525 EN Nijmegen, The Netherlands.

## Abstract

Convergent evidence from neuroimaging, computational, and clinical research has shown that the anterior temporal lobe (ATL) is critically involved in two key aspects of semantic cognition: the representation of semantic knowledge, and the executive regulation of this knowledge. Both are necessary for integrating features to understand concepts, and to integrate concepts to understand discourse. Here, we tested the hypothesis that these differential aspects of integration map onto different patterns of ATL connectivity. Specifically, we hypothesized that there are two overlapping modes of functional connectivity of the ATL that each predict distinct aspects of semantic cognition on an individual level. We used a novel analytical approach (connectopic mapping) to identify the first two dominant modes connection topographies (i.e. maps of spatially varying connectivity) in the ATL in 766 participants (Human Connectome Project), and summarized these into 16 parameters that reflect inter-individual differences in their functional organization. If these connection topographies reflect the ATL’s functional multiplicity, then we would expect to find a dissociation where one mode (but not the other) correlates with cross-modal matching of verbal and visual information (picture vocabulary naming), and the other (but not the former) correlates with how quickly and accurately relevant semantic information is retrieved (story comprehension). Our analysis revealed a gradient of spatially varying connectivity along the inferior-superior axis, and secondly, an anterior to posterior gradient. Multiple regression analyses revealed a double dissociation such that individual differences in the inferior-superior gradient are predictive of differences in story comprehension, whereas the anterior-posterior gradient maps onto differences in picture vocabulary naming, but not vice versa. These findings indicate that overlapping gradients of functional connectivity in the ATL are related to differential behaviors, which is important for understanding how its functional organization underlies its multiple functions.

## Introduction

Convergent evidence from neuroimaging, computational, and clinical research has shown that the anterior temporal lobe (ATL) is critically involved in two key aspects of semantic cognition: the representation of semantic knowledge, and the executive regulation of this knowledge (Lambon Ralph, Jefferies, Patterson, & Rogers, 2017). The first refers to the role of the ATL in representing concepts, which are distilled from rich, multi-modal, verbal and non-verbal experiences (see e.g., Martin & Chao, 2001; Patterson, Nestor, & Rogers, 2007; Rogers et al., 2004). They are not stored as unitary representations, but rather are widely distributed across modality-specific cortex, including somatosensory cortex, language areas, and affective brain areas. This idea resonates with early neuroscientific work (Meynert, 1866; Wernicke, 1874) that already argued that a web of connections forms the basis for the representation of concepts (Eggert, 1977; Lambon Ralph, 2014), and with more contemporary accounts of embodied cognition, which are supported by evidence that modality-specific information is activated in the corresponding brain areas when concepts are encountered (Mahon & Caramazza, 2008; Postle, McMahon, Ashton, Meredith, & de Zubicaray, 2008; Pulvermüller, 2005; Pulvermüller, Hauk, Nikulin, & Ilmoniemi, 2005).

Given that concepts are likely to be stored in a distributed fashion rather than in the form of unitary representations, it is necessary to integrate the distributed modality-specific features of a concept into a coherent representation, supported by “convergence zones” in the brain (Damasio, 1989). One popular theory is that the ATL plays a critical role in this integrative process by serving as a transmodal “hub” in which modality-specific information is mediated (Lambon Ralph et al., 2017). The so called “hub-and-spoke” model (Patterson et al., 2007; Rogers et al., 2004) suggests that the ATL serves as a highly interconnected hub, which receives input from the modality-specific “spokes”. Indeed, neuroimaging research has shown that the ATL receives input from regions involved in visual and auditory perception, as well as olfactory and limbic regions (Binney, Parker, & Lambon Ralph, 2012; Jackson, Hoffman, Pobric, & Lambon Ralph, 2016; Visser & Lambon Ralph, 2011).

Additional evidence for the role of the ATL as a hub in a “hub-and-spoke” configuration comes from the literature on semantic dementia (SD). In SD, relatively focal atrophy of the ATL (Schwartz, Marin, & Saffran, 1979; Snowden, Goulding, & Neary, 1989; Warrington, 1975) and its connectivity with modality-specific regions (Guo et al., 2013; Mummery et al., 1999) leads to impoverished semantic cognition, including lower performance on word naming, lexical fluency, and word reading tasks. These behavioral measures have been shown to correlate with resting cerebral glucose measures in the ATL, such that the degree of hypometabolism is related to the severity of the semantic impairment (Butler, Brambati, Miller, & Gorno-Tempini, 2009; Mion et al., 2010; Nestor, Fryer, & Hodges, 2006). This is further corroborated by post-mortem studies that have reported negative correlations between the amount of atrophy in the ATL and semantic scores (Mesulam et al., 2019; Mesulam, Thompson, Weintraub, & Rogalski, 2015; Yamamoto et al., 2009), and by meta-analytic work that has demonstrated a relationship between lesions in the left anterior temporal lobe and semantic naming errors (Mirman et al., 2015; Schwartz et al., 2009). Importantly, the semantic impairments observed in SD are largely independent of modality (Patterson et al., 2007), suggesting that the ATL plays a critical role in binding modality-specific information, rather than storing this information per se (Lambon Ralph, Sage, Jones, & Mayberry, 2010).

The literature on SD has been augmented with studies that established the criticality of regions for semantic representation using repetitive transcranial magnetic stimulation (rTMS) in non-patient populations. As expected, the application of rTMS to the ATL results in selective semantic impairments that mimic those observed in SD (Lambon Ralph, Pobric, & Jefferies, 2009; Pobric, Jefferies, & Ralph, 2007). In addition, rTMS application to the modality-specific “spoke” regions leads to category-specific deficits, whereas application to the ATL indeed results in category-general impairments (Pobric, Jefferies, & Lambon Ralph, 2010). Taken together, patterns of brain connectivity, impairments in SD, and rTMS-induced deficits all provide evidence for the idea that the ATL serves as a critical transmodal hub with functionally relevant and topographically organized connectivity with modality-specific regions (Guo et al., 2013; Patterson et al., 2007).

Semantic cognition also critically relies on the executive regulation of semantic knowledge: the (re)activation and selection of semantic information that is relevant for the task at hand, ongoing behavior, or current context, which itself is established through a mechanism of semantic control. For instance, the fact that pianos are heavy is not particularly relevant for playing the piano, but is highly relevant for moving it, which influences precisely which semantic aspects of “piano” should be activated (Moody & Gennari, 2010; Saffran, 2000). Previous work has suggested that the brain network that underlies semantic control is largely separate from the network for semantic representation (Lambon Ralph et al., 2017). Like the semantic representation network, the semantic control network is distributed: it consists of the lateral prefrontal cortex, posterior middle temporal lobe, intraparietal sulcus, pre-supplementary motor cortex, and the anterior cingulate-ventromedial prefrontal cortex, as well as the ATL hub (Chiou, Humphreys, Jung, & Lambon Ralph, 2018; Lambon Ralph et al., 2017).

Deficits in semantic control, for instance in semantic aphasia (SA) following stroke, manifest themselves as problems with semantic access (Lambon Ralph, 2014). In contrast with patients with SD, patients with SA have conceptual representations, but cannot access them (Lambon Ralph, 2014). For instance, patients with SA have difficulty naming pictures without any cues, but substantially improve when presented with a phonemic cue (for instance “oc” when naming “octopus”; Jefferies & Lambon Ralph, 2006; Jefferies, Patterson, & Ralph, 2008). This contrast between SD and SA suggests that having representations as well as being able to access to them are essential components of semantic cognition, and the dissociation found in lesion studies suggests that each might be supported by a distinct set of neural underpinnings (Jefferies & Lambon Ralph, 2006).

The ability to select and integrate context-relevant features of conceptual representations is essential for understanding and producing language in context, for instance in a sentence or a larger piece of discourse. Indeed, previous work has found that ATL activity increases when semantic information needs to be integrated across words in a sentence or across sentences in discourse (Brennan & Pylkkänen, 2017; Humphries, Love, Swinney, & Hickok, 2005; Stowe et al., 1998; Xu, Kemeny, Park, Frattali, & Braun, 2005). Tasks with high semantic control demands—such as pairing perceptually distinct but semantically similar concepts— evoke increased levels of activity in the ATL and semantic control system, and increased connectivity with traditional language areas such as inferior frontal gyrus (Chiou et al., 2018). Taken together, these findings suggest that semantic cognition relies on dynamic, task- and demand sensitive interactions between semantic representation and semantic control networks.

An open question, however, is how the neurobiological architecture of the ATL supports these dynamic interactions. One account is that the ATL exhibits “graded convergence” (Plaut, 2002), which refers to the idea that there are gradual shifts in function across the ATL. Indeed, accounts as early as Brodmann’s cytoarchitecture have suggested that transitions in the temporal cortex are graded rather than demarcated by sharp boundaries (Brodmann, 1909, in: Lambon Ralph et al., 2017). According to Plaut’s account (2002), gradual shifts in the ATL result from a topographic proximity bias such that areas of the ATL that are closer to modality-specific regions display higher degrees of specificity for that modality. Such a gradual, topographic organization is parsimonious as it allows for flexible interplay between modality-specific systems (Plaut, 2002).

Evidence from functional fMRI further corroborates the idea that the ATL’s semantic function is graded. In particular, modality-specificity appears to vary across the ATL, with function shifting from modality unspecific processing in ventrolateral ATL to modality specific processing in medial and superior regions (see Lambon Ralph et al., 2017 for an extensive review). For instance, task based fMRI work has shown a graded difference in ATL response to abstract, social, or modality-specific stimuli (Binney, Hoffman, & Lambon Ralph, 2016). Evidence from resting state fMRI functional connectivity analyses indeed has shown that ventrolateral ATL and anterior middle temporal gyrus are connected with regions involved in multimodal semantic processing, whereas superior temporal areas displayed connectivity with regions involved in auditory and language-related processing (Jackson et al., 2016; Murphy et al., 2017). Interestingly, these superior temporal areas (specifically, superior temporal gyrus) overlap with those that have been associated with combinatorial semantic processing (Hickok & Poeppel, 2007; Humphries et al., 2005; Lambon Ralph et al., 2017).

Jointly these findings support the notion that the ATL is embedded in multiple brain networks, including those that give rise to emergent semantic representations and semantic control networks. This necessitates that at least two overlapping modes of connectivity are embedded in the ATL, representing multiple axes of information convergence (Visser, Jefferies, Embleton, & Lambon Ralph, 2012) or function. Evidence from diffusion tensor imaging has indeed shown that termination points of white matter tracts are overlapping, suggesting a graded organization in terms of structural connectivity, potentially with multiple, overlapping connectivity profiles in a single region (Bajada, Haroon, et al., 2017; Bajada, Jackson, et al., 2017; Bajada, Trujillo-Barreto, Parker, Cloutman, & Lambon Ralph, 2019; Binney et al., 2012). However, most studies so far have assigned each termination point to a single white matter tract (but see Bajada, Jackson, et al., 2017), not taking into consideration that the ATL might contain multiple, overlapping modes of connectivity that simultaneously subserve implementation of representational and executive function.

The insight that the organization of the ATL might be best characterized in terms of multiple, overlapping connectivity profiles with gradual changes in function has fundamental implications for how ATL connectivity and its relationship with function should be analyzed and conceptualized. Analyses that treat parts or all of ATL as a spatially piecewise constant entity might not capture the true underlying organization of this region (Haak, Marquand, & Beckmann, 2018; Marquand, Haak, & Beckmann, 2017; Przeździk, Faber, Fernández, Beckmann, & Haak, 2019). Moreover, if indeed multiple overlapping modes of connectivity exist in the ATL, then failing to account for this multiplicity leads to biologically implausible results, as any unitary analyses would be conducted on the superposition of these modes (Haak et al., 2018; Jbabdi, Sotiropoulos, & Behrens, 2013; Kaas, 1997) which in itself might not follow biologically plausible trajectories

Here, we therefore take a novel approach to characterizing the functional organization of the ATL by decomposing its connectivity into overlapping modes of variation using Connectopic Mapping (Haak et al., 2018). This method is not only capable of capturing multiple, overlapping modes of spatially varying connectivity, but can also capture both gradual transitions as well as sharp boundaries within a region at the individual subject level. The ensuing connectopic maps—which represent the separate connection topographies at the individual level—can be summarized using a spatial statistics framework (Trend Surface Modeling; henceforth TSM) in order to facilitate statistical assessment, linking inter-individual variation in topographic connectivity to individual-level behavior.

We leveraged the wealth of the WU-Minn Human Connectome Project (HCP) data set (Van Essen et al., 2013, 2012) to establish the link between the organization of the ATL at the individual level—derived from resting state fMRI data using Connectopic Mapping—and ATL-mediated behavior. Specifically, the HCP data set contains two functional tasks that have previously been linked to activity and/or are affected by atrophy in the ATL (Binder, Swanson, Hammeke, & Sabsevitz, 2008; Mesulam et al., 2019; Rosazza et al., 2013), namely the NIH Toolbox Picture Vocabulary Test (TPVT; Gershon et al., 2013), and the Story Task (Binder et al., 2011). Importantly, although both tasks tap into semantic processing, the TPVT relies on cross-modal matching between an auditory presented word and a visually presented picture (i.e. multimodal integration of features to arrive at a meaning), whereas successful execution of the Story Task relies on combinatorial semantic processing (i.e. integration across meanings to understand discourse). As noted above, previous work has shown that similar ATL regions are active in both modality-specific processing and combinatorial semantic processing. If these represent two functional “modes” of the ATL, then we would expect that distinct, overlapping connection topographies underlie each behavior. To establish this, we derived the first two connection topographies in the ATL at the level of each individual participant, and subsequently linked each subject-specific gradient to individual differences in performance on each task.

A secondary open question is whether the link between the functional organization of the ATL and behavior is apparent in both hemispheres. Previous work has observed a leftward bias for word processing in the ventral ATL (e.g., Hoffman & Lambon Ralph, 2018; Rice, Ralph, & Hoffman, 2015). However, patients in whom the left ATL was resected still have (some) preservation of semantic knowledge (e.g., Bi et al., 2011; Rice, Caswell, Moore, Hoffman, & Lambon Ralph, 2018), suggesting that the right ATL also contributes to semantic cognition. We therefore established the brain-behavior relationships for each hemisphere separately.

## Methods

### Participants

We used the publicly available resting-state fMRI data from the WU-Minn Human Connectome Project S1200 release (Van Essen et al., 2012; see: https://db.humanconnectome.org). For the present study, we included all subjects who completed all of the four resting-state fMRI scans (r227 reconstruction version, *N* = 766 participants, 395 females, age range: 22-37). Because participants were drawn from families of twins and non-twin siblings, we accounted for the family structure of the HCP sample in the reported analyses using a permutation testing procedure implemented in FSL-PALM (Winkler, Ridgway, Webster, Smith, & Nichols, 2014).

### Data acquisition and preprocessing

The fMRI data of the HCP comprises multiband-accelerated (x8) scans with a temporal resolution of 0.72s and an isotropic spatial resolution of 2mm. Full details regarding the subject recruitment, ethical statements, and data acquisition protocols are described elsewhere (Smith et al., 2013; Van Essen et al., 2013). The resting-state data were preprocessed according to minimal processing pipeline procedures (Glasser et al., 2013), which included corrections for spatial distortions and head movements, registration to the T1w structural image, normalization to a common space using Multimodal Surface Matching procedures, global intensity normalization, high-pass temporal filtering using a cut-off of 2000 s, and rigorous denoising using ICA-FIX artifact removal procedures (FSL-FIX, Griffanti et al., 2014; Salimi-Khorshidi et al., 2014). The minimal processing pipeline included bringing the data in a standard grayordinate space and minimal smoothing (2mm) using a geodesic Gaussian surface smoothing algorithm (Glasser et al., 2013). No further spatial smoothing was applied.

### Connectopic mapping procedures

Connectopic mapping (Haak et al., 2018) is a data-driven approach for estimating connection topographies for regions of interest based on resting-state fMRI data. For each HCP subject, we derived the dominant and second-dominant modes of spatially varying connectivity (i.e. connectopic maps) in the left and right ATL separately, where the ATL was defined in each cerebral hemisphere as those regions of cortex encompassing the following cortical regions as provided by a multimodal atlas of human cortex: two regions in temporal polar cortex (TGd and TGv), two regions posterior to temporal polar cortex (TF and TE2a), one anterior middle temporal gyrus region (TE1a), two regions in the anterior superior temporal sulcus (STSva and STSda), the peri-entorhinal and ectorhinal complex (PeEc), and anterior auditory association area STGa on the superior temporal gyrus anterior to Heschl’s gyrus (Glasser et al., 2016). To this end, we used the connectopic mapping procedures described in previous work (Haak et al., 2018). Briefly, for each ATL vertex (gray-ordinate) we obtained a connectivity ‘fingerprint’ by correlating its time-series with SVD-transformed data from all vertices outside the ATL. These fingerprints were used to construct a vertex-wise similarity matrix using the eta-squared statistic, which was submitted to a locality-preserving non-linear manifold learning algorithm (Laplacian Eigenmaps; see Haak et al., (2018) for parameter details). The resulting eigenvectors convey connectopic maps and are ordered according to their eigenvalues (Belkin & Niyogi, 2003). In the present work, the two dominant connectopic maps were selected for further analysis.

### Trend surface modeling

To link individual differences in connectopic organization to behavior, we summarized each individual’s connection topography in terms of a limited number of spatial model parameters using trend surface modeling (Haak et al., 2018). This approach, which originally stems from geo-statistics, can be used to capture the overall spatial pattern of a (connectopic) map in a small number of coefficients by fitting a set of basis functions. We used polynomial basis functions defined in a spherical space and fitted using Bayesian linear regression, which also yields likelihood estimates of the model given the data. Starting estimation with a first-degree polynomial (a straight line with a slope), we added eight polynomials (2^nd^ – 8^th^ order) to the trend surface model. This was the optimal number of polynomials according to the Bayesian Information Criterion (BIC), which we computed from the model likelihoods for model selection purposes (testing model orders 1 through 10; in some cases, the optimal model order was slightly lower, but to ensure comparability whilst adequately accommodating the most complex topographies, we elected to take the 8^th^ order models forward for further analysis). The fitted polynomials yield 16 parameters that describe the connectopic organization of the surface of the ATL (eight polynomial basis functions on a two-dimensional surface).

### Behavioral data

We tested whether the ensuing trend surface parameters are meaningfully related to individual-level performance on two tasks: picture vocabulary naming (Gershon et al., 2013) and story comprehension (Binder et al., 2011), which are included in the National Institutes of Health Toolbox measures and the task-fMRI battery of the Human Connectome Project, respectively. In picture vocabulary naming task, participants were presented with an audio recording of a word and four pictures, and were asked to select that picture that most closely matched the meaning of the auditorily presented word. This test was designed for participants aged 3-85, and takes approximately four minutes to complete. We used the age-adjusted scale score (this score compares the score of the test-taker to others of the same age) as the dependent variable for the picture vocabulary task in subsequent analyses. For story comprehension, participants were presented with audio recordings of four brief stories (5-9 sentences, adapted from Aesop’s fables), followed by visually presented comprehension questions (2-alternative forced choice question about the topic of the story; for example, after a story about an eagle that saves a man who had done him a favor, participants were asked, *‘That was about revenge or reciprocity?’*; Barch et al., 2013; Binder et al., 2011). The stories were interleaved with four blocks of math trials that were auditorily presented, which were also followed by a visually presented 2-alternative forced choice question. Each participant listened to four stories, and performed four blocks of math tasks. In total, this task took approximately four minutes, which each block taking approximately 30 seconds. For both tasks, we took proportion of correct items divided by the median response time as a dependent measure for subsequent analyses. This measure captures variations in how quickly and accurately participants answered the questions.

## Results

### Dominant connectopic maps

We used Connectopic Mapping to reveal the first two modes of spatially varying connectivity in the ATL. We found that in both hemispheres, the dominant mode of variation runs along the inferior-superior axis, and the second along the anterior-posterior axis. An individual-level average projection of these gradients in the left hemisphere for an example subject can be found in Fig. 1. To visualize the full extent of these gradients along the ATL surface, we projected each gradient onto a sphere (middle panel Fig. 1). As expected, both connection topographies display a relatively gradual organization, with smooth rather than sharp transitions between ATL subfields.

**Figure 1.**
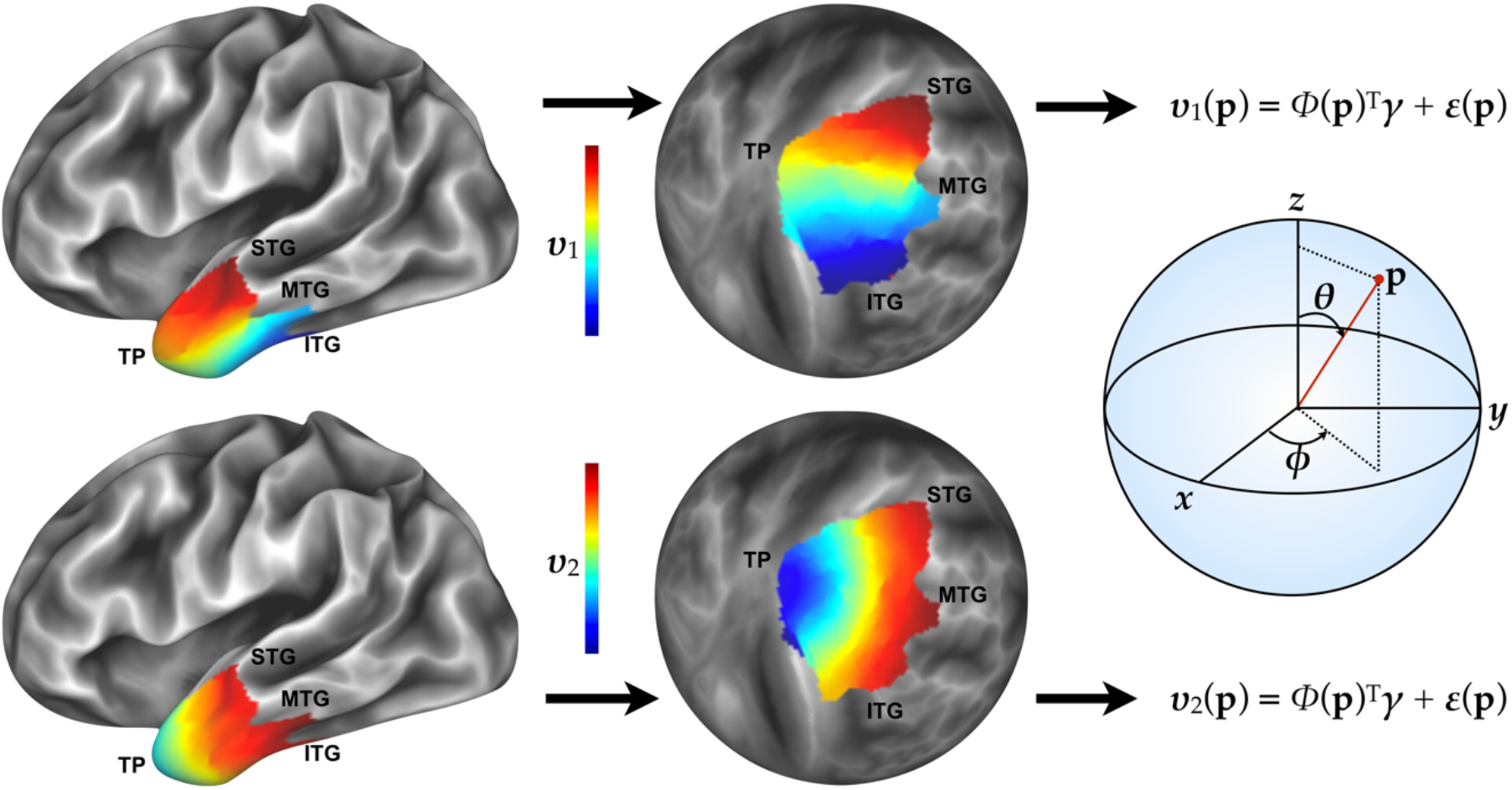
Overlapping gradients of functional connectivity in the anterior temporal lobe. Left: the dominant modes of spatially varying connectivity in the ATL are situated along the inferior-superior (top) and anterior-posterior axes. Shown is an example gradient at the individual subject-level. Similar colors indicate similar connectivity ‘fingerprints’. Middle and right: To enable statistical inference over these gradient maps, they were projected into spherical space where we applied trend-surface modelling (using spherical coordinates) to capture the spatial structure with a limited number of model coefficients. TP = Temporal Pole, STG = Superior Temporal Gyrus, MTG = Middle Temporal Gyrus, ITG = Inferior Temporal Gyrus.

### Behavioral associations

For each individual, we used trend surface modeling (see above and Fig. 1) to summarize their connectopic maps in terms of a low number of spatial model parameters. We entered these parameters (16) as explanatory variables in a multiple regression framework to estimate the relationship between individual-level connectopic organization and individual-level behavior. In these analyses, we controlled for head motion (i.e. mean frame-wise displacement) and age, by computing the partial *R*^2^ as (RSS_covariates_ - RSS_parameters+covariates_) / RSS_covariates_. Subjects with poor trend surface model fits were excluded from this analysis by applying the interquartile rule for outlier detection to the variance explained by the model (first gradient: LH = 66, RH = 80; second gradient: LH = 82, RH = 57). Analyses were conducted for each hemisphere and each behavioral variable separately, and accounted for the family structure in the data.

For the left hemisphere, we found that individual differences in the inferior-superior gradient were significantly related to differences in story comprehension (partial *R*^2^ = .06, *p* = .001). Individual differences in the organization of the anterior-posterior gradient displayed a significant relationship with differences in picture vocabulary naming (partial *R*^2^ = .04, *p* = .02). Importantly, the organization of the first gradient did not significantly explain picture vocabulary naming (partial *R*^2^ = .02, *p* = .79), and conversely, the second gradient’s organization was not significantly related to story comprehension (partial *R*^2^ = .03, *p* = .17) (Fig. 2a). To address the specificity of these relationships for language processing, we assessed the relationship with performance on the math control task, and found no significant relationships with either gradient.

**Figure 2.**
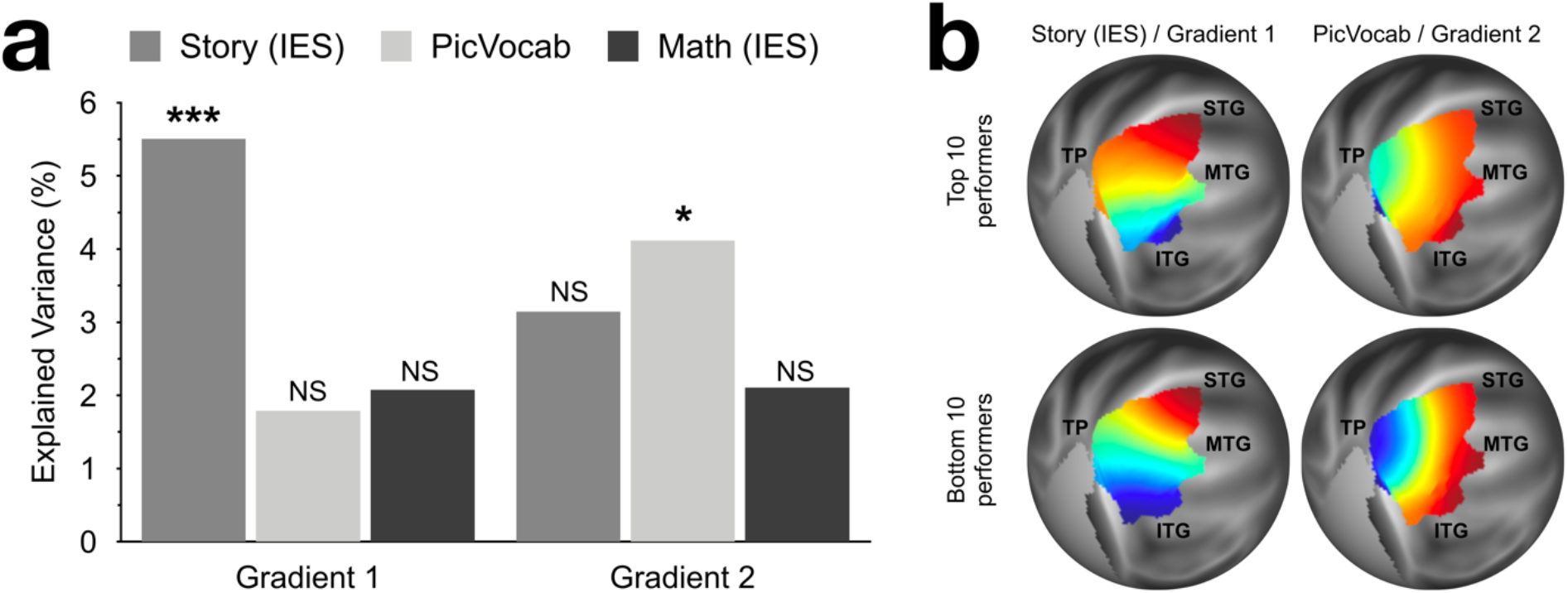
Associations between the left ATL connection topography gradients and behavior. **a.** Linear regression analyses show that individual differences in the trend surface parameters that describe the inferior-superior gradient are associated with differences in story comprehension, but not in picture vocabulary naming or math. Vice versa, individual differences in the anterior-posterior gradient are related to differences in picture vocabulary naming, but not story comprehension or math. **b.** Average ATL connection topography gradients of the top and bottom 10 performers on the story comprehension and picture vocabulary tasks.

For the right hemisphere, we found no significant associations between the inferior-superior gradient and any of the tasks (story comprehension: partial *R*^2^ = .03, *p* = .19; picture vocabulary naming: partial *R*^2^ = .02, *p* = .56; math: partial *R*^2^ = .02, *p* = .68). The anterior-posterior gradient similarly displayed no significant relationships with any of the tasks (story comprehension: partial *R*^2^ = .01, *p* = .97; picture vocabulary naming: partial *R*^2^ = .03, *p* = .25; math: partial *R*^2^ = .02, *p* = .61). This observation suggests that the relationship between connectopic organization and language-related semantic cognition might be left-lateralized. This is in line with earlier work that suggests that there is a leftward bias for word processing in the ATL (Hoffman & Lambon Ralph, 2018; Rice, Ralph, & Hoffman, 2015), and is in line with the observation that language function is predominantly left-lateralized in the majority of the healthy population (Springer et al., 1999).

An important concern for all studies into ATL pertains to potential susceptibility-induced signal loss, resulting in a temporal signal-to-noise ratio (tSNR) reduction in inferior ATL that was also present in the HCP data. Consequently, we observed a strong correlation between the overall spatial distribution of the temporal signal-to-noise ratio (tSNR) and the dominant (i.e. superior-inferior) gradient in both hemispheres (LH: *r* = .63; RH: *r* = .65). However, individual differences in the dominant gradient were only very weakly related to individual differences in the tSNR maps (LH: *R*^2^ = .001; RH: *R*^2^ = .002; comparison between the cross-subject covariances of the tSNR and gradient maps), and unlike the left inferior-superior gradient, trend-surface modeling of the tSNR maps did not yield a significant association with story comprehension (partial *R*^2^ = .03, *p* = 0.29). Thus, the tSNR reduction in inferior ATL appears to be unrelated to the individual differences in the dominant ATL gradient that explain ATL mediated behaviors.

### Individual differences and cortical projections

To illustrate how inter-individual variations in connectopic organization (i.e. the distribution of functional connectivity across the ATL) underlie differences in behavior, we visualized the average gradient for the top and bottom 10 performers on the story comprehension and picture vocabulary naming tasks (Fig. 2B). The gradients are scaled from 0 to 1 and color coded along that range, so that similar colors represent that the vertices in the ATL have similar extrinsic connectivity profile. Different colors likewise denote maximally different extrinsic connectivity. Thus, it can be gleaned from the transition from red to blue in each gradient that there are clear inter-individual differences in organization in both gradients.

To additionally illustrate these differences in terms of connectivity with the rest of cortex, we visualized the connectivity profiles of the extremes of each gradient (Fig. 3). This was achieved by performing a whole brain seed-based connectivity analysis using the most anterior, posterior, inferior and superior ATL subfields (i.e. TGd, TE2a, PeEc and STSda, respectively; Glasser et al., 2016) as seeds. The mean time-series was extracted for each seed, and correlated against the vertex-wise time-series across the rest of the brain while controlling for the mean time-series in the other three seed regions. Fig. 3 shows the average connectivity strength (i.e. partial correlation) across 20 unrelated subjects. Note that this analysis reveals the unique connectivity of the extreme ends of the two gradients, but due to its seed-based nature does no longer account for functional multiplicity within the individual subfields. The results of this analysis are therefore likely an over-simplification of the true connectivity patterns and should be taken as illustration only.

**Figure 3.**
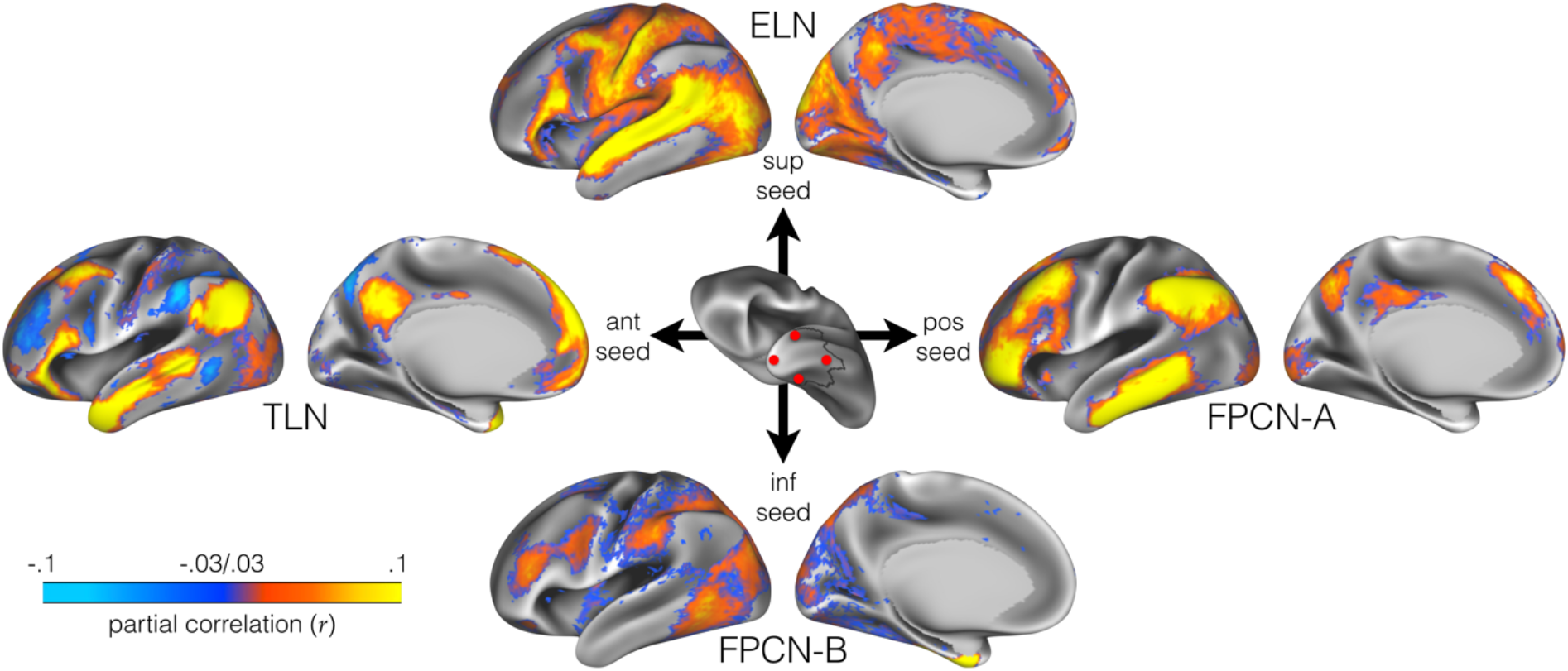
Unique seed-based functional connectivity patterns associated with the extremes of each gradient. See main text for details. Maps were thresholded at *z_r_* ≥ 2. The superior-inferior ATL gradient appears to be underpinned by connectivity moving gradually from the extended language network (ELN) and modality-specific brain regions at the superior extreme, to frontoparietal control network (FPCN) subsystem B at the inferior extreme. The anterior-posterior gradient appears to follow a gradual transition of connectivity with traditional language regions (TLN) at the anterior extreme, to frontoparietal control network (FPCN) subsystem A at the posterior extreme.

The superior extreme of the superior-inferior gradient shows strong positive correlations with traditional language areas (BA44/45), precuneus, sensory-motor cortices, visual, and auditory areas. The inferior extreme, on the other hand, displays strong connectivity with dorsolateral prefrontal cortex, the inferior parietal lobule, and extrastriate occipital cortex. The connectivity patterns associated with the inferior extreme resemble frontoparietal control subsystem B (Dixon et al., 2018). Differences visualized in Fig. 2B suggest that the better performers on the story task might have relatively more neural resources allocated to receiving input from the modality-specific regions (relatively large red area, corresponding to the superior connectivity profile) compared to the poor performers (smaller red area).

For the anterior extreme of the anterior-posterior gradient, we observed strong positive correlations with traditional language areas (BA44/45/47), lateral parietal cortex, lateral occipital areas, and posterior cingulate cortex, and negative correlations with dorsolateral prefrontal cortex, the inferior parietal lobule, and posterior middle temporal cortex. In contrast, the posterior extreme of this gradient displays only strong positive correlations, and thus connectivity, with dorsolateral prefrontal cortex and the inferior parietal lobule, plus correlations with the insula, language areas, visual areas (occipital pole), posterior cingulate cortex, and precuneus. These connectivity patterns resemble frontoparietal control subsystem A (Dixon et al., 2018). Better performers appear to have relatively more neural resources allocated to connectivity with the latter constellation of networks (relatively large red area, corresponding to the posterior connectivity profile) compared to poor performers.

## Discussion

We tested the idea that the functional organization of the ATL can be meaningfully described in terms of multiple functionally relevant, overlapping gradients of connectopic organization. We used a novel data analytical approach, connectopic mapping, to derive these maps at the individual subject level, and found two overlapping modes of organization: a primary gradient along the superior-inferior axis, and a secondary gradient along the anterior-posterior axis. We established their functional meaningfulness through correlations with individual level behavior, and found a double dissociation such that individual differences in the superior-inferior gradient are related to story comprehension, whereas individual differences in the anterior-posterior gradient map onto differences in picture vocabulary naming. These differences in connectopic organization likely reflect differences in connectivity with modality-specific brain regions, language areas, and control regions: compared with poor performers, better performers might have more neural resources allocated to interactions with the extended language network as well as modality specific networks (in support of story comprehension) or frontoparietal control subsystem A (in support of picture vocabulary naming).

This study is the first to link interindividual differences in ATL organization to differences in ATL-mediated behaviors in a population-based sample. Our findings are in line with previous work has shown that the integrity of the ATL and its connectivity—which can be negatively affected by hypometabolism and atrophy—are critically related to semantic cognition in clinical populations. Importantly, the current work shows that not only adverse conditions but also interindividual differences in organization can be related to semantic cognition. These findings are in line with other recent work that has shown that the fine-grained connectopic organization of other brain regions, such as the human striatum and hippocampus, is predictive of individual level goal-related behavior and recollection memory, respectively (Marquand et al., 2017; Przeździk et al., 2019), suggesting that the characterization of brain organization in terms of these connectopic maps is a fruitful avenue for gaining a better understanding of brain-behavior relationships.

Here, we focused on the relationship between ATL organization and semantic cognition, limited to language comprehension (picture vocabulary task, story comprehension task). We found meaningful relationships between the organization of the left—but not the right—ATL and these behaviors, suggesting that functional relationships with language comprehension are left lateralized. This observation is corroborated by experimental and meta-analytical work that has shown that indeed, there is a leftward bias for word processing in the ventral ATL (Hoffman & Lambon Ralph, 2018; Rice, Ralph, & Hoffman, 2015). This is not to say, however, that there is no contribution of the right ATL to language processing, as there is a wealth of literature showing contributions of both left and right ATL (e.g., Bi et al., 2011; Rice, Caswell, Moore, Hoffman, & Lambon Ralph, 2018; Rice, Hoffman, & Lambon Ralph, 2015). It seems plausible that the left laterality that we observed here is driven by connectivity with the traditional language network (see Fig. 3), given its relative left laterality in the majority of the population. Furthermore, it is possible that inter-subject variability in other, non-verbal semantic tasks, such as picture matching (Rice, Caswell, Moore, Lambon Ralph, & Hoffman, 2018), is related to differences in connectopic organization in the bilateral ATLs. Future research could scrutinize these relationships further to increase our understanding of how the functional organization of both left and right ATL underlies behavior.

What do the results tell us about the relationship between the functional organization of the left ATL and its functional role in picture vocabulary matching and story comprehension, and in semantic cognition more generally? One interesting observation is that each gradient appears to be connected with the frontoparietal control network, also known as the multiple demand system, which is thought of as a distributed, domain general control system (Duncan, 2010). Recent work, however, has suggested that there are two distinct subsystems in the frontoparietal control network that display differential connectivity with the dorsal attention network and default mode network of the brain (Dixon et al., 2018). In particular, one subnetwork (FPCN_A_) is thought to be preferentially coupled with the default mode network, and has meta-analytically been linked to processes such as decision-making, mentalizing, and emotional processing. Importantly, its preferential connectivity with the default mode network suggests that it might regulate the conceptual-associative knowledge supplied by the default mode network. The second subnetwork (FPCN_B_), on the other hand, is most strongly connected to the dorsal attention network, and has been linked to reading, semantic processing, action, and attention. The role of the FPCN_B_ in these processes might be to exert executive control over the activation or maintenance of mental representations that are relevant to the task at hand.

Interestingly, there is substantial overlap between FPCN_A_ and FPCN_B_ on the one hand and the connectivity profiles of the posterior extreme of the anterior-posterior gradient and the inferior extreme of the superior-inferior gradient on the other hand. This architecture might facilitate semantic cognition by providing integration or convergence between conceptual content and control: the fact that the anterior-posterior gradient appears to run from connectivity with traditional language regions and the default mode network to FPCN_A_ might underpin its role in picture vocabulary matching by supplying conceptual content (anteriorly) and exerting control over that content (posteriorly). Similarly, the superior-inferior gradient runs from connectivity with the extended language network and modality-specific regions to FPCN_B_, which might exert executive control over the modality-specific and linguistic information necessary for the task at hand, suggesting a potential role in combinatorial language processing.

These findings resonate with the more widespread idea that brain networks are organized in a hierarchical gradient of processing (Dixon et al., 2018; Dixon, Fox, & Christoff, 2014; Huntenburg, Bazin, & Margulies, 2018; Margulies et al., 2016; Murphy et al., 2018). Recent work has proposed that at the macroscale, brain areas are organized along a global gradient that runs from sensorimotor (i.e. unimodal) regions to transmodal (i.e. heteromodal) regions (Dixon et al., 2018; Huntenburg et al., 2018). Our findings suggest that to some extent, the same principle might govern the organization of the ATL: we observed two gradients that each run from conceptual processing (either linguistic or conceptual content, or sensorimotor information) to domain general control networks that might regulate this input. Such an intrinsic organization might be optimal for integrating information across different scales: integration across features to make sense of concepts, and across concepts to make sense of discourse. Given the topographical organization of other regions within the networks in which the ATL is embedded (Dixon et al., 2018, 2014; Rebecca L Jackson, Bajada, Lambon Ralph, & Cloutman, 2019; Przezdzik, Haak, Beckmann & Bartsch, 2019), an open question is how the ATL fits in the larger hierarchy of the cortical gradient(s).

In a broader topographic context, the ATL gradual organization seem to be in line with the topographical organization of the traditional language areas. Previous connectopic mapping work showed a connectivity gradient in the left inferior frontal gyrus (IFG) (Przezdzik, Haak, Beckmann, & Bartsch, 2019), which in accordance with the Memory, Unification and Control model (Hagoort, 2005) may reflect the organization of the unification of syntax, semantic and phonology components, as well as the memory and control processes in the IFG. Importantly, the cortical projections of this IFG gradient seems to follow the anterior-posterior trajectory in the temporal lobe, which resemble the anterior-posterior organization in the ATL, important for the representation of the semantic knowledge and the task that relies on the cross-modal matching of verbal and visual information. This overlap in the temporal lobe may therefore suggest broader topographic organization of language processing, characterized by similar connectivity patterns of multiple language areas (here IFG and ATL) in the temporal cortex.

Our results also agree with previous work that has suggested multiple gradients of information convergence in the temporal lobe. Indeed, region of interest analyses have shown lateral (superior temporal gyrus → inferior/middle temporal gyrus ← fusiform gyrus) and longitudinal (toward ATL) axes of convergence in the temporal lobe (Visser et al., 2012), which have recently been corroborated by data-driven functional parcellations of temporal cortex (Jackson, Bajada, Rice, Cloutman, & Lambon Ralph, 2018). In the present work, we showed that such an organization exists not only at the level of the temporal lobe as a whole, but also within the anterior temporal lobe and we were able to link these modes of organization to individual differences in semantic cognition.

What aspect of the underlying neurobiology drives the gradient-like organization that we observe here? Given the structural connectivity of the ATL with modality-specific and language areas, it is likely that the gradients are in part driven by terminations of long-range white matter tracts. In particular, recent work has shown that there are white matter tracts that terminate along the superior temporal gyrus-inferior temporal gyrus axis (Bajada, Jackson, et al., 2017; Blazquez Freches et al., 2020). These in part overlap with terminations of the arcuate fasciculus along the posterior-anterior axis (Bajada, Haroon, et al., 2017; Blazquez Freches et al., 2020). It is therefore possible that the resting-state fMRI derived connectopic maps that we obtained reflect the individual-level organization of white matter tract terminations in the ATL. However, ATL is also known to have strong within-ATL connectivity that is not captured with diffusion-weighted imaging. Although variation in the superior and posterior extremes might be driven by white matter tract terminations, gradual variations in adjacent areas might reflect local microstructure (Mollink et al., 2019; Vos de Wael et al., 2018). Further research could shed light on the relationship between white matter macro- and microstructure and connectivity measures derived with functional MRI.

Our work has focused on the two dominant modes of spatially varying connectivity in the ATL, demonstrating that these gradients bear functional relevance at the individual level. However, this is not to say that the ATL comprises exactly two overlapping modes of functional organization. It is possible that additional gradients capture other (known or unknown) organizational principles of the ATL, such as its graded specialization for modality-specific semantics (Binney et al., 2016; Visser et al., 2012), further reflecting its hub-and-spoke-like organization. How many meaningful, superimposed gradients are present in the ATL is thus far an open question. This also implies that the analysis of unique connectivity of the extreme ends of the two gradients presented here likely is an over-simplification, as it may not fully account for this functional multiplicity.

In conclusion, we showed that the functional organization of the ATL can be described in terms of overlapping gradients of spatially varying connectivity, and that the two dominant modes are meaningfully related to individual level behavior. These findings are not only important for understanding the functional role of the ATL in semantic cognition, but also have implications for the conceptualization and study of its organization: its architecture should be considered within the context of its embedding in multiple brain networks that are associated with conceptual processing and cognitive control, which in turn are likely to be organized in terms of gradients. Our findings suggest that organization of the ATL facilitates its role in semantic cognition by uniting modality-specific information, such as features with concepts, linguistic information, and cognitive control, facilitating both the integration of features into concepts, and integration across concepts to make sense of discourse.

## Acknowledgments

This work was supported by the Netherlands Organization for Scientific Research Vidi Grant No. 864-12-003 (to CFB), Veni Grant No. 016.Veni.171.068 (to KVH), Veni Grant No. VI.Veni.191G.001 (to MF), NWO-CAS Grant No. 012-200-013 (to CFB, providing funds to MF), and Gravitation Programme Grant No. 024.001.006 (providing funds to IP). Data were provided by the Human Connectome Project, WU-Minn Consortium (Principal Investigators: David Van Essen and Kamil Ugurbil; 1U54MH091657) funded by the 16 NIH Institutes and Centers that support the NIH Blueprint for Neuroscience Research; and by the McDonnell Center for Systems Neuroscience at Washington University.

## Notes

### Competing Interest Statement

The authors have declared no competing interest.

